# Reconstitution of lipid nanoparticle-mediated siRNA transfer across endosomal membrane mimics

**DOI:** 10.64898/2026.11.02.763664

**Authors:** A. Spadea, A. Singh, L. Redlingshöfer, S. Sorrentino, Y. Kalaidzidis, M. Zerial

**Affiliations:** Max Planck Institute of Molecular Cell Biology and Genetics (MPI-CBG), Dresden, Germany; Human Technopole Foundation (HT), Milan, Italy

## Abstract

Endosomal escape limits RNA delivery by lipid nanoparticles (LNPs). If escape requires membrane rupture, productive delivery would be intrinsically coupled to membrane damage and its potential toxicity. Here, we establish a cell-free system based on giant unilamellar vesicles (GUVs) with an early endosome-like lipid composition to dissect the mechanism of siRNA transfer and determine whether it can occur while membrane barrier function is preserved. Acidification promoted LNP-membrane association, lipid redistribution and siRNA translocation into the receiving lumen without detectable dissipation of the transmembrane pH gradient. Complementary cryo-electron tomography revealed local interfacial lipid reorganisation without detectable membrane rupture. Fluorescence correlation spectroscopy revealed heterogeneous diffusion of translocated siRNA, consistent with free and lipid-associated species. Pre-formed RNA-lipid complexes exhibited pH-dependent dissociation and heterogeneous diffusion, supporting lipid association as a possible origin of slower luminal diffusion. Serum or apolipoprotein-E preincubation of LNPs reduced siRNA translocation into the GUV lumen. Together, these findings support a model where siRNA-lipid association facilitates membrane crossing without sustained loss of barrier function. This platform provides a means to dissect LNP-mediated RNA delivery and develop formulations with improved efficacy while limiting membrane damage.

## INTRODUCTION

Lipid nanoparticles (LNPs) have become a leading platform for RNA therapeutics, including the first FDA-approved siRNA drug patisiran, and the mRNA vaccines against COVID-19 (Du Rietz et al. 2020; Buschmann et al. 2021). LNP-mediated delivery rests on the ability of ionizable lipids to remain near-neutral in circulation and become protonated inside the acidifying endosomes, promoting interactions with the endosomal membranes and release of the RNA cargo into the cytosol. Nevertheless, only a small fraction of internalized RNA reaches the cytosol, with escape efficiency estimated in the range of only 1-8%, depending on formulation, cellular context and measurement method (Gilleron et al. 2013; Paramasivam et al. 2021; Wittrup et al. 2015; Jozić et al. 2026; Sabnis et al. 2018; Patel et al. 2020). Endosomal escape therefore remains the central bottleneck limiting the potency, and hence the required dose, of RNA-LNP therapeutics (Dowdy et al. 2022). Understanding how RNA crosses the endosomal membrane is therefore central to improving LNP-mediated delivery.

Endosomes are dynamic, compositionally distinct compartments. Early endosomes have a luminal pH ∼6.3-6.8, and mature into late endosomes with more acidic pH ∼5-5.5 and the ability to fuse with lysosomes. Where along this trajectory escape occurs is debated, with evidence pointing to early endosomal/recycling compartments (Gilleron et al. 2013; Paramasivam et al. 2021), although a later, more acidic endosomal stage has also been proposed (Sahay et al. 2013; Wittrup et al. 2015). Mechanistically, the prevailing model invokes protonated ionizable lipids ion-pairing with anionic endosomal lipids to form non-bilayer, hexagonal-phase structures that locally destabilize the membrane (Semple et al. 2010; Hafez, Maurer, and Cullis 2001). This is distinct from bona fide membrane fusion, such as the back-fusion of intraluminal vesicles with the limiting endosomal membrane (van der Goot and Gruenberg 2006). Because RNA-loaded LNPs have an internal organisation different from conventional bilayer vesicles, the relationship between their interfacial lipid rearrangements and classical vesicle membrane fusion remains to be established.

Live-cell imaging studies have also reported endosomal membrane damage and leakage following LNP treatment (Johansson et al. 2025; Omo-Lamai et al. 2025). The extent of damage and its relationship to cargo delivery can depend on formulation and exposure conditions (Munson et al. 2021; Bates et al. 2025). RNA-LNP therapeutics operate within a narrow margin between the dose required for efficacy and the dose at which cytotoxicity and innate immune activation become dose-limiting (Hou et al. 2021). At sufficiently high intracellular loads, endosomal damage can likely be induced, but this may not reflect the mechanism operating under physiologically and therapeutically relevant conditions (Munson et al. 2021). Endosomal membrane damage can activate inflammatory pathways (Omo-Lamai et al. 2025). If productive RNA delivery obligatorily requires membrane rupture, increasing escape efficiency would intrinsically couple therapeutic activity to a potentially harmful event, constraining efforts to improve efficacy without increasing toxicity. Establishing whether RNA can cross the membrane while preserving compartmentalisation is therefore central to identifying delivery mechanisms that could separate these outcomes.

Several questions therefore remain open. Does RNA translocation require sustained membrane rupture or can it occur while compartmentalisation is preserved (Semple et al. 2010; Hafez, Maurer, and Cullis 2001)? How does RNA transfer relate to LNP association and lipid redistribution at the membrane? What physical states does RNA adopt after crossing, and how do proteins encountered in biological fluids affect these events?

Addressing these questions in cells is challenging because uptake, trafficking, membrane composition and acidification are tightly coupled, making their individual contributions difficult to dissect experimentally. Moreover, endosomal escape events are rare and stochastic, and the nanoscale membrane rearrangements underlying RNA translocation occur below the spatial resolution of conventional light microscopy. Galectin recruitment reports exposure of normally luminal glycans following endosomal membrane damage, but does not directly resolve the transient membrane discontinuities or local nanoscale lipid rearrangements that may enable RNA translocation (Aits et al. 2015). Reporter release and knockdown provide cellular readouts of cargo delivery and biological activity. Even when single-molecule imaging localises individual escape events (Paramasivam et al. 2021), the molecular interactions and nanoscale membrane rearrangements at the precise site of RNA translocation remain unresolved.

Cell-free approaches, including bulk liposome assays such as FRET-based lipid mixing and dye leakage (Zheng et al. 2023), supported lipid bilayers (Aliakbarinodehi et al. 2022; Aliakbarinodehi et al. 2024), and lipid monolayers (Spadea et al. 2022), have resolved important aspects of LNP-membrane interactions. A complementary requirement is to measure cargo transfer into a separate receiving compartment while assessing membrane barrier function and the physical state of the transferred RNA.

Here, we address these questions using a cell-free system based on giant unilamellar vesicles (GUVs) with defined early endosome lipid composition (Cezanne et al. 2020) and topology. In this system, an acidic, LNP-containing external medium, representing the endosomal lumen, is separated from a neutral vesicle interior that serves as the receiving compartment for RNA translocation, representing the cytosolic side of the membrane. Combining fluorescence imaging, intravesicular pH measurements, Cryo-electron tomography and fluorescence correlation spectroscopy (FCS), we distinguish membrane association and lipid redistribution from siRNA transfer into the receiving compartment and characterise the diffusion behaviour of the transferred cargo.

## RESULTS

### Acidification triggers LNP-membrane association and lipid redistribution

Endosomal acidification is a key environmental cue encountered by LNPs after cellular uptake, promoting interactions between ionizable lipids and endosomal membranes. Dissecting the ensuing events in cells is challenging both because uptake, trafficking and escape are interdependent and because the small dimensions of endosomes limit access to the membrane interface where RNA translocation occurs. Although single-molecule imaging can localise individual escape events, resolving the underlying molecular rearrangements while achieving sufficient throughput and independently varying experimental conditions remains difficult. To combine optical accessibility with control over membrane composition and pH, we established a reconstituted system based on giant unilamellar vesicles (GUVs) with a lipid composition mimicking early endosomal membranes (Cezanne et al. 2020). Their large size allows membrane-associated signals to be spatially distinguished from cargo in the receiving lumen, enabling complementary measurements of LNP-membrane interactions and RNA transfer under defined conditions. In this system, the GUV lumen serves as a receiving compartment corresponding to the cytosolic side, while the external bulk medium corresponds to the endosomal lumen. GUVs are formed at pH 7.2, which therefore defines the pH of their lumen, whereas the pH of the external medium can be adjusted independently by adding an acidic buffer, thereby establishing a pH gradient across the membrane comparable to that experienced by an LNP-containing endosome and the cytosol (Fig. 1a). We incubated GUVs for 10 min with MC3-based LNPs carrying Alexa488-labelled siRNA and DiD-labelled lipids (Fig. 1b), with the external pH set to 6.5, representative of the early endosomal lumen, or kept at 7.2 as a neutral control, allowing us to follow RNA and lipid behaviour simultaneously at the LNP-membrane interface.

**Figure 1.**
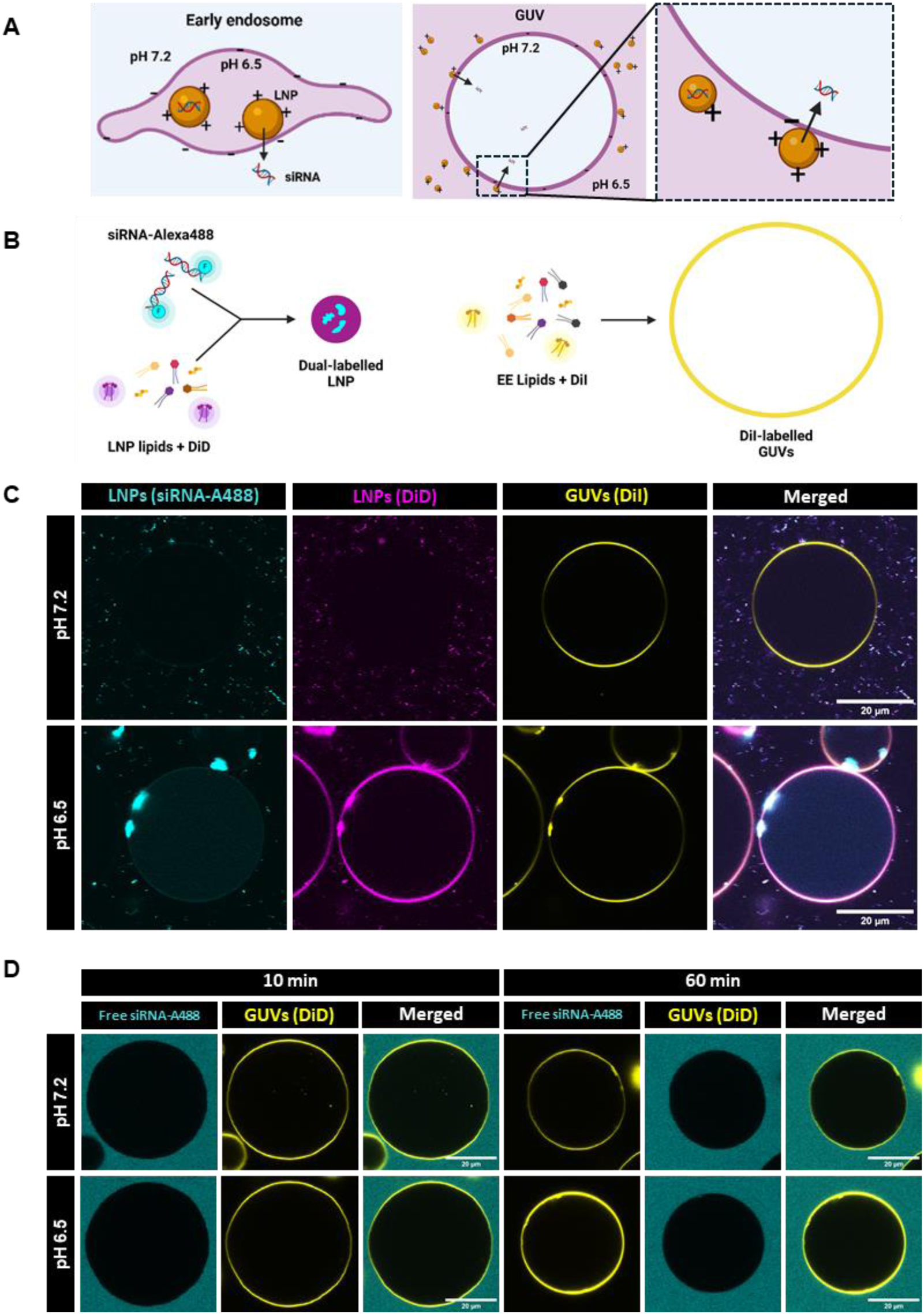
Acidification triggers LNP-membrane interaction and lipid redistribution in early endosome-mimicking GUVs. **a,** Schematic of the reconstituted system. GUVs mimicking the lipid composition of early endosomal membranes are formed at pH 7.2, which defines the pH of the GUV lumen (cytosol equivalent). The pH of the external medium (endosomal lumen equivalent) is adjusted by addition of an acidic buffer. **b,** Schematic of the dual-labelling strategy: siRNA-Alexa488 and LNP lipids (DiD) are co-formulated into dual-labelled LNPs, while GUVs are independently labelled with DiI during membrane preparation. **c,** Representative confocal images of GUVs incubated with dual-labelled LNPs (125 nM siRNA-Alexa488, cyan; DiD, magenta) for 10 min at an external pH of 7.2 (top) or 6.5 (bottom), with the GUV membrane labelled with DiI (yellow). **d,** Representative confocal images of GUVs incubated with free siRNA-Alexa488 (no LNPs) at an external pH of 7.2 (top) or 6.5 (bottom), for 10 min or 60 min. In the absence of an LNP carrier, siRNA shows no detectable association with the GUV membrane or lumen at either pH or time point, confirming that the membrane-associated clusters and luminal signal in c depend on LNP-mediated delivery. Scale bars, 20 µm.

At pH 7.2, we detected no appreciable association between LNPs and the GUV membrane (Fig. 1c). In contrast, at pH 6.5 LNPs associated readily with GUVs, accumulating as discrete siRNA-positive clusters at the outer membrane surface (Fig. 1c; see Supplementary Fig. S2a for representative full fields of view). We also observed diffuse siRNA fluorescence in the vesicle lumen, consistent with translocation of RNA across the membrane (Fig. 1c). Free siRNA, in the absence of LNPs, showed no association with GUVs at either pH (Fig. 1d; Supplementary Fig. S2b) up to 60 min, confirming that RNA accumulation at the membrane and in the lumen depends on the LNP carrier. This was confirmed quantitatively by measuring the fluorescence photon count rate from a confocal detection volume (the same point-detection setup used for FCS measurements, see Fig. 4 and Methods), which showed abundant free siRNA in the bulk solution surrounding GUVs but only background-level signal within the GUV lumen at either pH (Supplementary Fig. S1).

The DiD signal, tracking LNP-associated lipid dye, displayed a different spatial pattern from the RNA. Whereas siRNA formed discrete membrane-associated clusters and a diffuse luminal signal, DiD became progressively more homogeneously distributed along the GUV membrane at pH 6.5, consistent with transfer and lateral redistribution on the vesicle bilayer of lipid-associated dye (Fig. 1c). These distinct spatial readouts allow membrane association, lipid redistribution and luminal RNA accumulation to be assessed separately, suggesting that lipid redistribution and RNA release into the lumen can be distinguished experimentally. Together, these data show that acidification alone is sufficient to trigger LNP interaction with an early endosome-like membrane, accompanied by lipid rearrangements and partitioning of siRNA between membrane-associated and luminal pools.

### LNP-mediated siRNA translocation occurs without detectable loss of the transmembrane pH gradient

The observation of RNA in the GUV lumen prompted us to examine whether translocation is accompanied by disruption of the membrane. One way to approach this problem is to determine whether the receiving compartment retained its pH. If the integrity of the GUV bilayer were compromised following the LNP interaction, protons would be expected to flow down their gradient, acidifying the GUV lumen. We therefore used SNARF-5, a ratiometric pH-sensitive dye, to measure pH inside and outside the vesicles. Following external acidification, the luminal pH remained at ∼7.2 at 10 and 60 min, both in the absence and presence of LNPs (Fig. 2a). Measurements of bulk pH confirmed external acidification in each condition (Fig. 2b), confirming that the stable luminal readout reflects genuine membrane behaviour rather than inconsistent acidification of the external medium.

**Figure 2.**
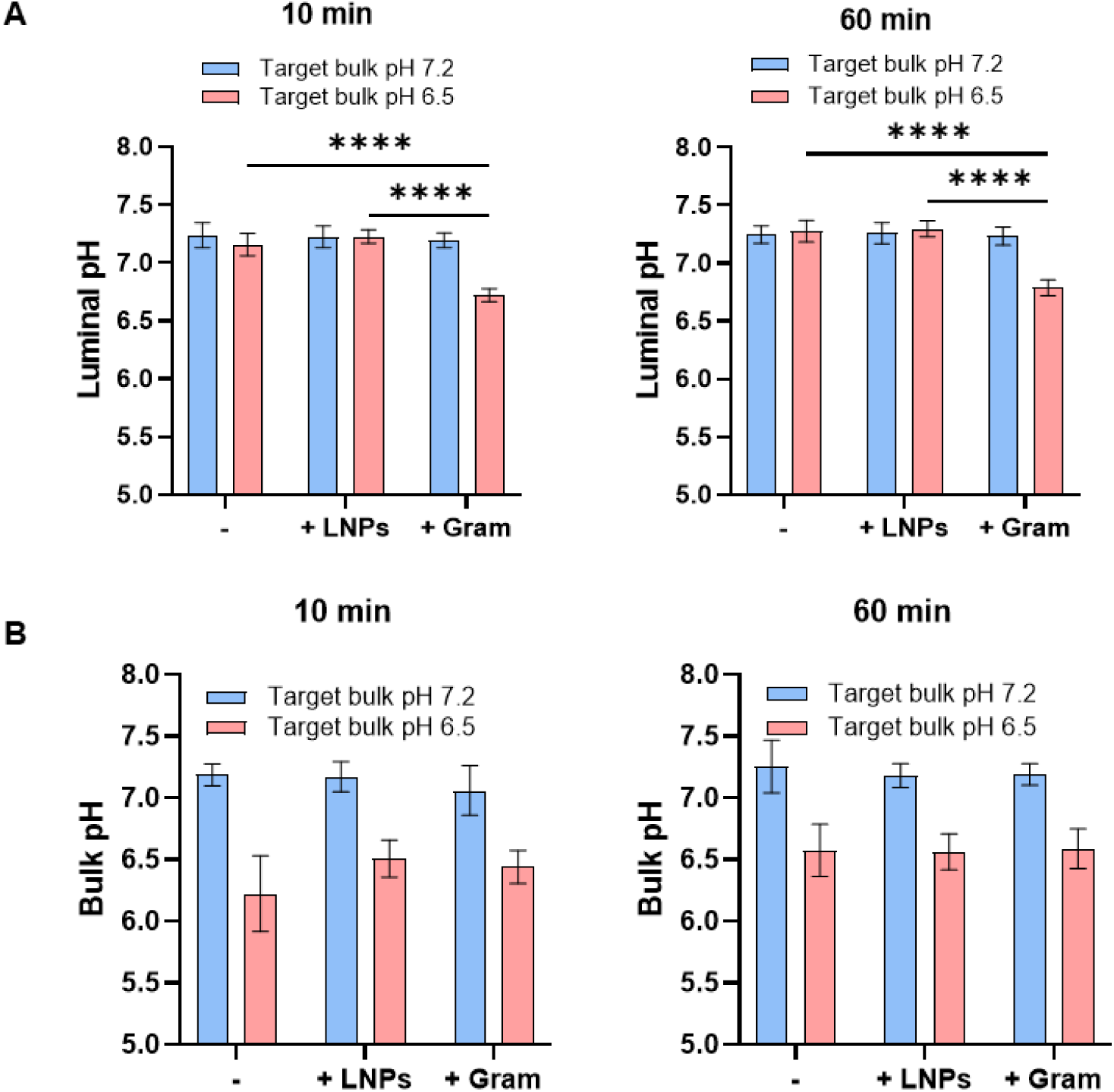
LNP-mediated siRNA translocation occurs without detectable loss of the transmembrane pH gradient. **a,** Luminal pH of GUVs at neutral pH or after external acidification to pH 6.5, measured in the absence of LNPs (-), in the presence of LNPs (+LNPs), or in the presence of the pore-forming peptide Gramicidin (+Gram) as a positive control for membrane permeabilization, at 10 min (left) and 60 min (right). **b,** Corresponding measured bulk (external) pH for the same conditions and time points, documenting the external pH achieved in each condition. Data in a and b are median ± s.d. from ≥20 GUVs (a) or 3 bulk measurements per image across 10 images (b) per condition, from N = 3 independent experiments. Statistical comparisons were performed by two-way ANOVA with Tukey’s multiple comparisons test; only significant comparisons are shown. ****P < 0.0001.

As a positive control, we used the selective cation channel forming peptide gramicidin A (Gram), which forms channels that permit proton flux across lipid bilayers (Akeson and Deamer 1991; Sun et al. 2020). When Gram-treated GUVs were exposed to external pH 6.5, luminal pH dropped to ∼6.7 at 10 min and remained low at 60 min (Fig. 2a). This control demonstrates the sensitivity of the assay to detect membrane permeabilization to protons under the assay conditions. The bulk pH readout was, as expected, unaffected by the presence of LNPs or gramicidin at either pH (Fig. 2b), confirming that gramicidin acts specifically on the GUV membrane rather than altering the buffer itself. The absence of comparable acidification in LNP-treated GUVs indicates retention of the pH gradient, but does not exclude transient and spatially restricted membrane defects.

Together, these results show that LNP treatment did not cause equilibration of protons between the two compartments under the experimental conditions tested, suggesting that membrane integrity was not compromised.

### LNP-membrane interaction induces local lipid rearrangement, visualized by cryo-electron microscopy

The preservation of a proton gradient suggests that the endosomal mimic membrane cannot present ruptures upon LNP-GUV interaction. To examine the structural changes that accompany LNP-membrane interaction, and whether they involve open holes or ruptures, we used cryo-electron tomography. Because cryo-electron microscopy requires vitrification in an ice layer only ∼100-300 nm thick for electron transparency, we incubated LNPs with large unilamellar vesicles (LUVs) of the same early endosome-like lipid composition as the GUVs, as a size-compatible membrane substrate. The smaller vesicles are compatible with the thin vitreous ice required for transmission imaging and provide an accessible substrate for resolving the membrane-LNP interface (Fig. 3).

**Figure 3.**
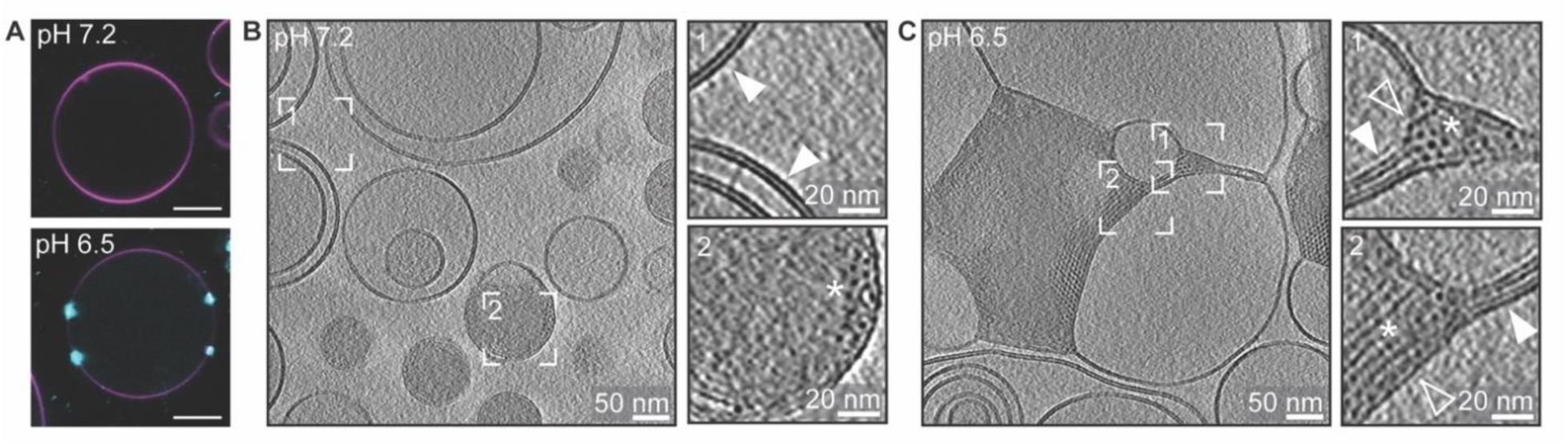
LNP-membrane interaction induces local lipid rearrangement, visualized by cryo-electron tomography. **a,** Representative confocal overview images of a GUV incubated with LNPs, showing membrane-associated siRNA-positive LNP clusters at pH 6.5 and no LNP-membrane interactions at pH 7.2. **b** and **c,** Z-slices through a reconstructed cryo-electron tomography volume of LUVs of the same early endosome-like lipid composition as in **a**, incubated with LNPs at pH 7.2 (**b**) or pH 6.5 (**c**) for 10 min. Boxed regions correspond to magnified areas on the right, highlighting intact LUV bilayer membranes (filled arrowheads), and LUV membrane remodelled at contacts with LNP clusters (empty arrowheads). Ordered phases of LNP lipids in the inert single LNPs (**b**, **box 2**) and membrane-interacting LNP clusters (**c**, **box 1** and **2**) are marked by an asterisk. Scale bar, 10 µm (**a**), 50 nm (**b**, **c**), 20 nm (**insets**).

In the complementary LUVs preparation at pH 7.2, LNPs remained as discrete, individual particles, with LUV membranes intact and separate from the LNPs (Fig. 3b). At pH 6.5, LNPs instead formed assemblies in which individual particle boundaries became indistinct and LUV membranes directly continuous with the LNP clusters, with no resolvable gap between bilayer and LNP material (Fig. 3c). At these contact sites, the bilayer was replaced by regularly spaced striations and honeycomb arrays of dots (Fig. 3c, boxes 1 and 2). Orthogonal views of such domains showed a dotted lattice in one plane and parallel striations in the two orthogonal planes (Supplementary Fig. S4), the signature of lipid cylinders packed on a lattice, i.e. an inverted hexagonal (HII)-type arrangement, and not of stacked bilayers, which would not produce a dotted lattice in any section. Notably, similar ordered lamellar and hexagonal arrangements, marked by an asterisk, were also detected within individual, membrane-distal LNPs (Fig. 3b, box 2), indicating that this internal organisation is an intrinsic feature of the particle itself rather than one induced solely by membrane contact. Together, these observations support pH-dependent, local structural reorganisation at LNP-membrane contacts, through which the bilayer’s native architecture is locally lost without evidence of discrete membrane discontinuities connecting the two aqueous compartments.

### Translocated siRNA exhibits heterogeneous diffusion behaviour

The accumulation of siRNA in GUVs with preservation of the pH gradient and membrane continuity at the ultrastructural level raises the question of how this hydrophilic, negatively charged molecule crosses the membrane. One possibility is that association with ionizable lipids partially shields the RNA’s charge and increases its lipophilicity, facilitating passage through the lipid bilayer. If these interactions persist after translocation into a neutral pH environment over time, the GUV’s lumen would be predicted to contain both free siRNA and siRNA-lipid assemblies. The two siRNA populations are expected to differ substantially in size and hence in diffusion coefficient. We therefore used fluorescence correlation spectroscopy (FCS), which resolves diffusing species by their diffusion coefficient at single-molecule sensitivity, to directly examining the diffusion behaviour of siRNA translocated into the GUV lumen (Fig. 4a). Because the diffusion coefficient scales inversely with the hydrodynamic radius, the small free siRNA-Alexa488 (a ∼13kDa duplex of a few nm) is expected to diffuse substantially faster than LNPs (∼100 nm in diameter) (Fig. 4a). As described in the Methods, we first determined the diffusion coefficient of free siRNA-Alexa488 in 7 mM HEPES buffer (∼116 µm²/s; Supplementary Fig. S5c), and then measured the same siRNA, together with siRNA-Alexa488 formulated in LNPs, in the GUV reaction buffer. In GUV reaction buffer, free siRNA-Alexa488 had a diffusion coefficient comparable to the HEPES reference within the experimental variation, and unchanged between external pH 7.2 and 6.5 (Fig. 4c). LNP-associated siRNA showed a similarly low diffusion coefficient (∼5-10 µm²/s) at both pH 7.2 and 6.5, indicating that LNPs remain intact as particles outside the GUV irrespective of external pH.

**Figure 4.**
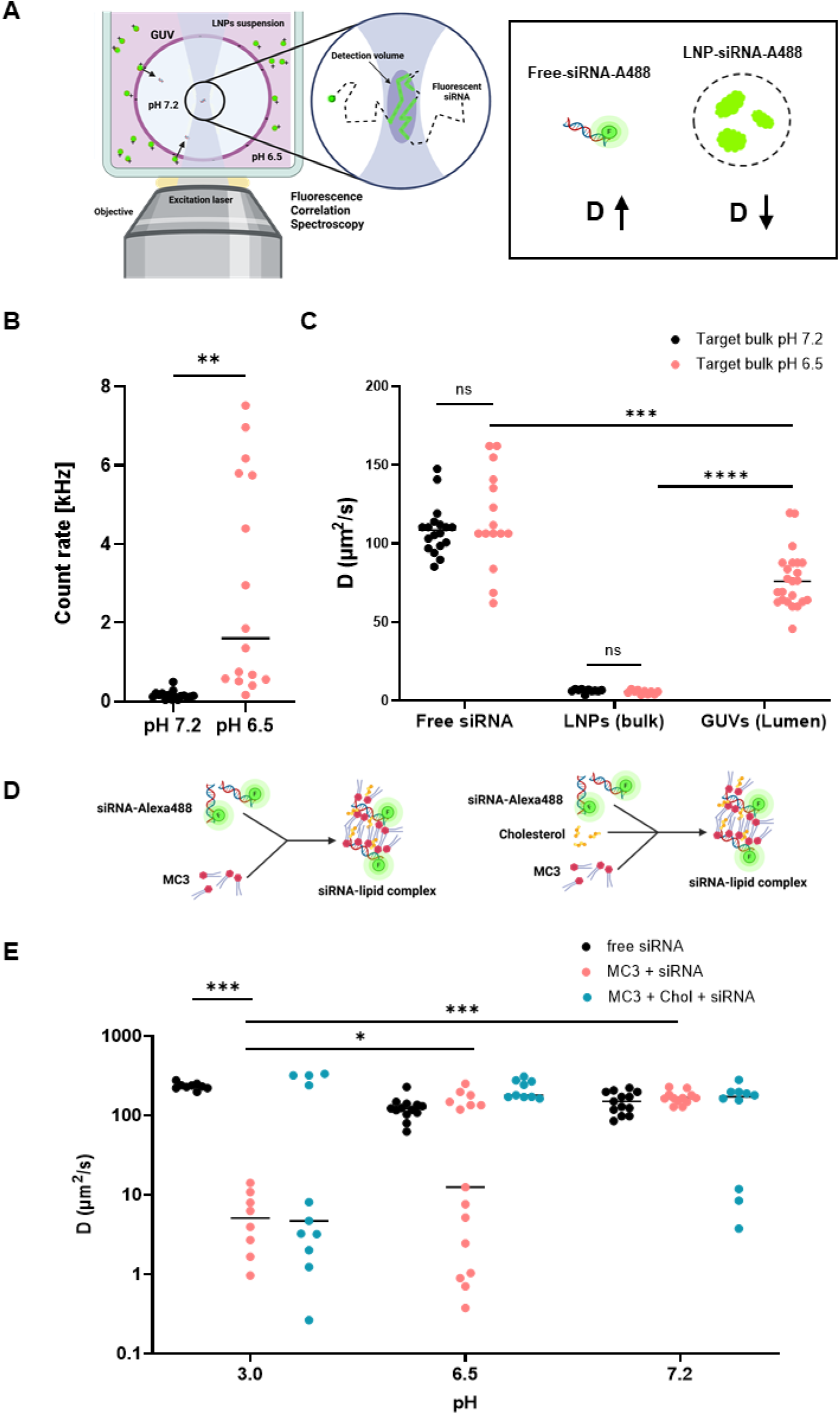
Transferred siRNA exhibits heterogeneous diffusion behaviour. **a,** Schematic of the FCS measurement: the confocal detection volume is positioned in the GUV lumen, 10 min after LNP addition and pH adjustment, to determine the diffusion coefficient (D) of translocated siRNA-Alexa488. Free siRNA diffuses faster (D↑) than siRNA encapsulated within LNPs (D↓), where multiple molecules diffuse together within a larger particle. **b,** Photon count rate measured inside the GUV lumen at pH 7.2 and pH 6.5 (data from 16 GUVs pooled together from 4 independent experiments, N=4). **P =0.0011. Statistical analysis unpaired t-test with Welch’s correction. **c,** Diffusion coefficients of free siRNA-Alexa488 (in HEPES and in GUV imaging buffer), of LNPs (in GUV imaging buffer, at pH 7.2 and pH 6.5), and of siRNA measured inside the GUV lumen at pH 6.5. All autocorrelation curves were fitted using maximum entropy method (MEM), which does not assume a fixed number of diffusing populations therefore all the detected multiple diffusion coefficients are shown together. No autocorrelatable signal was detected inside the GUV lumen at pH 7.2, therefore not shown. **d,** Schematics of the pre-formed siRNA-MC3 and siRNA-MC3-cholesterol complexes. **e,** Diffusion coefficients of free siRNA-Alexa488 and of pre-formed siRNA-MC3 and siRNA-MC3-cholesterol complexes (N/P 5) in citrate buffer at pH 3.0, 6.5 and 7.2. All autocorrelation curves were fitted as in c. Each point is an individual measurement, pooled from 3 or 4 independent experiments (N = 3 for all conditions except free siRNA at pH 6.5 and pH 7.2, N = 4); horizontal lines indicate the median of the pooled measurements. For visualization, all measurements are shown, whereas statistical analysis was performed on the average of each independent experiment, to avoid treating measurements from the same experiment as independent. Comparisons were made by one-way ANOVA assuming a lognormal distribution, with Tukey’s multiple comparisons test across all pairs; only comparisons addressing the pH-dependence of complex formation, and the difference from free siRNA at pH 3, are shown. *P < 0.05, ***P < 0.001.

We then positioned the confocal volume inside the GUV lumen, away from the membrane, to assess siRNA translocation, incubating GUVs with LNPs in the external medium at pH 7.2 or 6.5, as in Fig. 1. At external pH 7.2, luminal fluorescence was close to background (photon count rate 0.045 ± 0.006 kHz) (Fig. 4b), consistent with the absence of LNP-membrane interaction and consequent siRNA translocation at neutral pH (Fig. 1). In contrast, at pH 6.5, luminal fluorescence was readily detected in the GUV lumen (2.902 ± 2.716 kHz) (Fig. 4b), with apparent diffusion coefficients spanning a broad range, approximately 50-120 µm²/s (Fig. 4c). The fastest values approached the free siRNA reference, whereas the slower values were consistent with larger RNA-containing assemblies, but away from the diffusion coefficient of intact LNPs. This distribution demonstrates heterogeneous diffusion behaviour across luminal recordings. The slower diffusing species could be siRNAs remaining partially associated with lipids.

To test whether RNA-lipid association could account for slow diffusion, we prepared siRNA-MC3 and siRNA-MC3-cholesterol complexes at pH 3, omitting the helper phospholipid and PEG-lipid of the complete LNP formulation, and measured diffusion after sequential shifts to pH 6.5 and 7.2 (Fig. 4d and e). At pH 3, siRNA-MC3 complexes diffused significantly more slowly than free siRNA (∼5 µm²/s). Their diffusion coefficient increased significantly when pH was brought to 6.5 and further at pH 7.2, where values approached the free-siRNA reference, consistent with a pH-dependent loss of RNA-lipid association. At pH 6.5, individual measurements included both free-siRNA-like and slower values, indicating partial dissociation. siRNA-MC3-cholesterol complexes also showed slow diffusion at pH 3 (median ∼5 µm²/s), but neither the difference from free siRNA nor the changes with pH reached significance, because complex formation at pH 3 varied between independent experiments (in one experiment no slow species was detected). We therefore could not resolve an effect of cholesterol on complex formation or on its pH-dependence.

Together, these measurements showed heterogeneous diffusion behaviour of transferred siRNA, ranging from free-siRNA-like to substantially slower values. The complementary lipid-siRNA complex experiments support persistent siRNA-lipid association as possible origin of the slower luminal species. Following membrane transfer, the shift from an acidic to neutral environment could lead to dissociation of siRNA from lipid-associated to fully free.

### Serum pre-incubation and ApoE reduce LNP-membrane interactions and luminal cargo accumulation

Proteins encountered in biological fluids can alter LNP interactions with membranes, forming a protein corona (Francia et al. 2020), and influence positively and/or negatively uptake and endosomal escape. For example, adsorption of ApoE on LNPs influences positively their uptake into hepatocytes via LDL-receptor mediated endocytosis (Akinc et al. 2010; Pattipeiluhu et al. 2022) and can restructure the LNP shell and core (Sebastiani et al. 2021). On the other hand, protein corona has been proposed to have a negative effect on endosomal escape (Voke et al. 2025; Aliakbarinodehi et al. 2022). To test whether the GUV platform recapitulates aspects of this physiological modulation, particularly endosomal escape, we first compared untreated LNPs with LNPs pre-incubated with fetal bovine serum (FBS) to mimic the adsorption of a complex, physiologically relevant set of proteins onto LNPs as in circulation. To then dissect the contribution of a single, established protein species, we focused on ApoE, one of the best-studied LNP corona proteins. To assess its effect in this system, we titrated ApoE during preincubation and assessed LNP-GUV membrane interaction and RNA translocation using the same FCS-based readouts (count rate, luminal siRNA concentration and diffusion coefficient) established in Fig. 4.

The siRNA fluorescent signal differed in appearance at the membrane: naked LNPs formed few, large clusters, whereas LNPs pre-incubated with FBS appeared as numerous, smaller puncta distributed along the membrane (Fig. 5a). The latter produced also a much lower luminal siRNA signal than the untreated control (Fig. 5a). The decrease of luminal signal prevents reliable estimation of luminal concentration (C) and diffusion coefficient (D). Therefore, we report photon count rate which does not require autocorrelation fitting and could be measured for all screened GUVs. In the case of the ApoE pre-incubation, the signal was high enough for FCS fitting criteria. Therefore, we performed titration of ApoE concentration to further dissect how this pre-incubation affects the physical state of translocated RNA.

**Figure 5.**
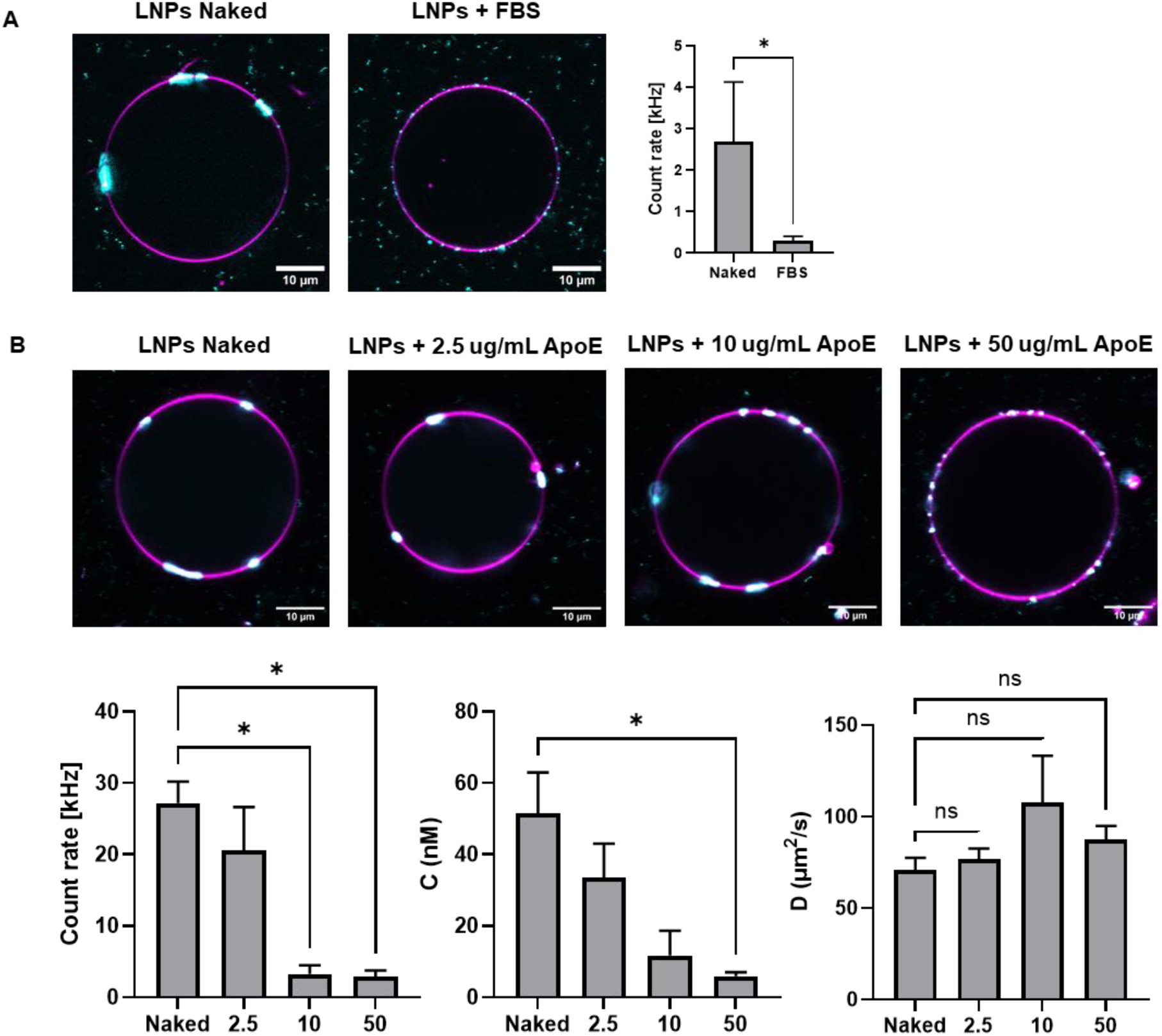
Serum pre-incubation and ApoE reduce LNP-membrane association and luminal siRNA accumulation. **a,** Representative confocal images of GUVs (magenta, DiD) incubated with naked LNPs or LNPs pre-incubated with FBS corona (cyan, siRNA-Alexa488). All conditions were imaged with identical acquisition settings, and images are displayed with identical display ranges. Right, count rate measured by FCS inside the GUV lumen for naked and FBS-corona LNPs. Bars show the mean ± s.e.m. of per-experiment medians across N = 3 independent experiments (naked: n = 35 GUVs screened; FBS: n = 35 GUVs screened). Count rate was log-transformed prior to a paired two-tailed t-test (assuming a lognormal distribution). *P < 0.05. Luminal concentration (C) and diffusion coefficient (D) are not reported for this condition: the low luminal signal resulting from reduced translocation meant that only the 28.6% of FBS-corona GUVs met the FCS fitting quality-control criteria (see Methods) — insufficient for statistical comparison. **b,** Representative confocal images of GUVs incubated with naked LNPs or LNPs pre-incubated with 2.5, 10 or 50 µg/mL ApoE. Below, count rate, C and D as in a, for naked LNPs and increasing ApoE concentrations. Bars show the mean ± s.e.m. of per-experiment medians across N = 3 independent experiments (for all conditions ≥75% of screened GUVs met the FCS fitting quality-control criteria). Count rate and C were log-transformed prior to repeated-measures one-way ANOVA (assuming a lognormal distribution) with Dunnett’s multiple comparisons test against naked LNPs; D was compared on linear values by repeated-measures one-way ANOVA with Dunnett’s multiple comparisons test. *P < 0.05 vs naked; ns, not significant.

Increasing ApoE concentrations progressively shifted the membrane-associated LNP pattern from large clusters towards numerous smaller puncta, resembling the FBS condition (Fig. 5b). ApoE pre-incubation reduced also luminal fluorescence count rate in a concentration-dependent manner, with significant differences at 10 and 50 µg/mL relative to untreated LNPs. The fitted apparent concentration was reduced significantly at 50 µg/mL (Fig. 5b). The diffusion coefficient of luminal siRNA remained comparable across all ApoE concentrations, with no statistically significant change detected among analysable luminal recordings.

Together, these results show that serum and ApoE pre-incubation reduce LNP clustering at the membrane and luminal siRNA accumulation, consistent with reports of corona-mediated inhibition of LNP delivery (Voke et al. 2025), with no significant change in apparent diffusion of the translocated siRNA species. The system provides a means to investigate how protein exposure affects membrane-crossing activity independently of cellular uptake, although the contributions of particle restructuring, protein binding and free protein require further distinction.

## DISCUSSION

Understanding how RNA crosses the endosomal membrane requires resolving how LNPs interact with the membrane, redistribute their lipids and induce local membrane rearrangements that enable a hydrophilic, highly charged cargo to traverse the hydrophobic bilayer interior. Here, we established a platform that resolves these processes across a free-standing early-endosome-like GUV membrane separating two aqueous compartments. The system recapitulates acidification-dependent interactions of ionizable lipid-containing LNPs with the membrane and translocation of siRNA in the receiving GUV lumen, representing the cytosolic side. Acidification promoted LNP-membrane association, lipid redistribution and luminal siRNA accumulation without detectable equilibration of the transmembrane pH gradient. Complementary cryo-electron tomography of LUVs and LNPs at low pH revealed local membrane remodelling without detectable open ruptures or holes on the membrane separating the aqueous compartments. These findings support the idea that siRNA can be transferred across a lipid bilayer with an early-endosome-like composition through local lipid rearrangements without sustained loss of membrane barrier function.

A key question in the field of RNA therapeutics delivery is how a highly charged RNA molecule can traverse the membrane while its overall barrier function is preserved. LNPs are densely packed structures with a lipid and cargo rich core (Chen 2026; Sebastiani et al. 2021). They lack the conventional bilayer architecture of vesicles and, thus, cannot be assumed to follow the classical bilayer-bilayer fusion such as that of intraluminal vesicles and exosomes fusing with the limiting membrane of endosomes/multi-vesicular bodies (van der Goot and Gruenberg 2006). Cryo-electron tomography revealed pronounced reorganisation at the LNP-membrane interface. Whereas LNPs remained discrete at neutral pH, acidic conditions produced assemblies with indistinct particle boundaries, in which the bilayer was replaced by regularly spaced striations and honeycomb arrays. Orthogonal views identified these as hexagonally packed cylinders of an inverted hexagonal (HII)-type phase. This is consistent with the mechanism proposed by Hafez, Maurer and Cullis, in which interactions between cationic lipids and anionic membrane phospholipids promote HII formation (Hafez, Maurer, and Cullis 2001). In this phase, lipid monolayers surround aqueous channels, with polar headgroups facing the channels and hydrocarbon chains directed outwards. Disordered inverse hexagonal organisation has also been described within mRNA-containing LNP cores (Yanez Arteta et al. 2018), raising the possibility that related arrangements arise during LNP-membrane interaction.

We propose that association with ionizable lipids facilitates siRNA transfer by partially shielding its charge and coupling local lipid reorganisation to cargo translocation (Fig. 6). In this model, membrane crossing and complete dissociation of siRNA from lipids are distinct steps. Exposure to the neutral pH receiving compartment would favour lipid deprotonation and weaken electrostatic interactions with RNA, allowing dissociation following translocation. Consistent with this model, luminal siRNA exhibited heterogeneous diffusion behaviour, ranging from free-siRNA-like values to substantially slower values compatible with larger assemblies. Pre-formed siRNA-lipid complexes likewise showed slow diffusion, with neutral pH favouring free-siRNA-like diffusion. These findings support siRNA-lipid complexes as a possible explanation for slower luminal diffusion. Interestingly, recovery of both mRNA and ionizable lipid in endosome-derived extracellular vesicles also suggests that lipid association can persist during intracellular processing (Maugeri et al. 2019). This distinction matters when interpreting escape efficiency: RNA reaching the receiving compartment may differ in its degree of lipid association and, therefore, its availability for downstream activity. Our platform measures both lipid redistribution and siRNA transfer into a separate receiving compartment, where the diffusion of transferred cargo can be characterised.

**Figure 6.**
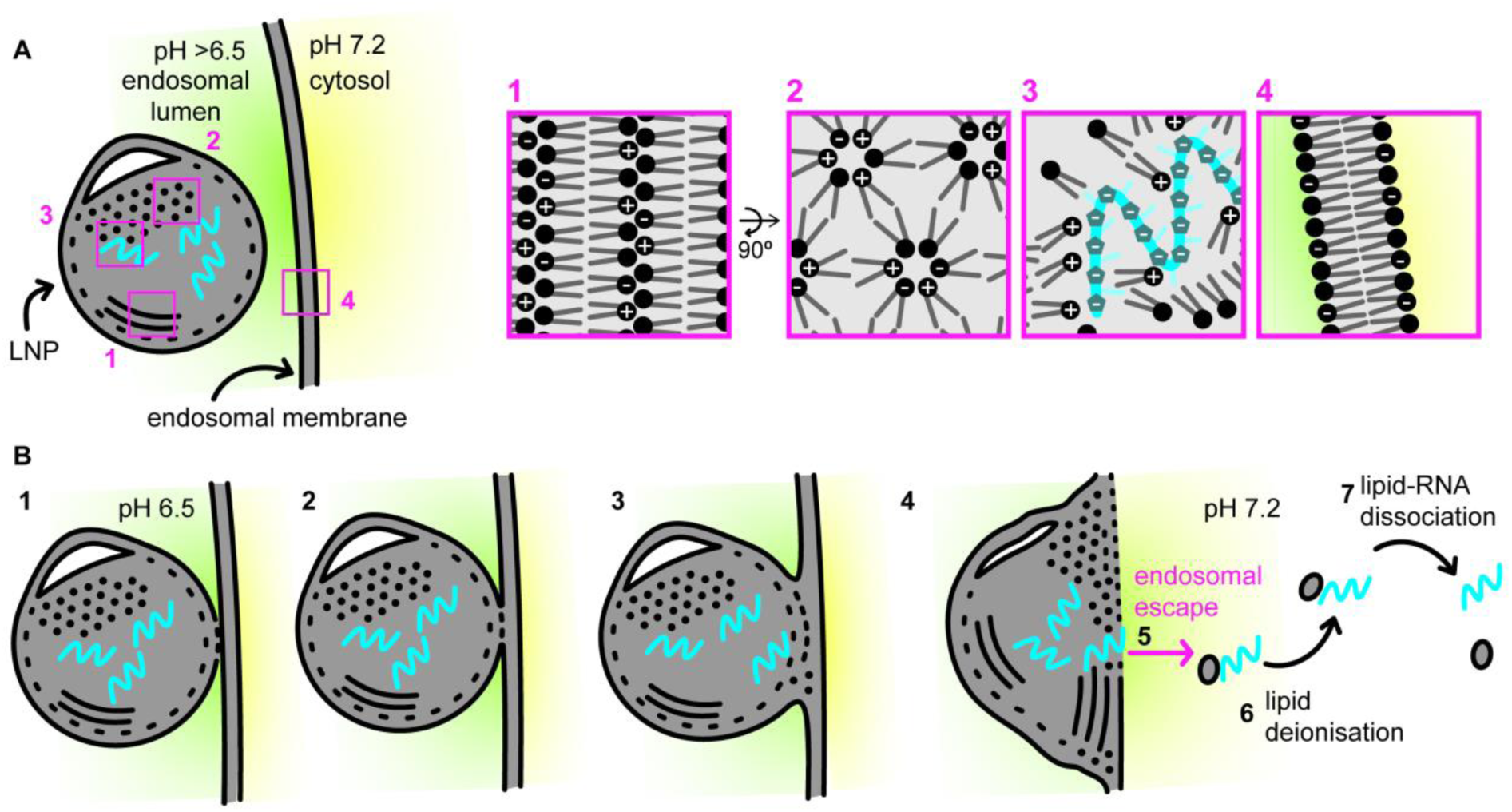
Proposed model of LNP-mediated and lipid-assisted siRNA transfer across an endosomal membrane. **a,** At neutral pH, LNPs remain discrete and show no association with the membrane. Boxes 1-4 illustrate the membrane bilayer, possible ordered lipid arrangements within LNPs and RNA-lipid association. The phospholipid bilayer of the endosome is negatively charged (**box 1**). While lipids within LNP are mostly in a disordered phase, they may locally arrange in an ordered fashion (**box 2**, hexagonal phase top view and **box 3**, hexagonal phase side view) and act as a solvent for the encapsulated RNA (**box 4**). **b**, Upon endosomal acidification (1), the ionizable lipids become protonated, promoting LNP-membrane association and lipid interspersion with the inner leaflet of the endosomal bilayer (2). LNPs then mix further with the endosomal membrane (3) leading to rearrangement of the second bilayer leaflet (4) towards hexagonal arrangement as shown in A (**box 2** and **3**). Without an intact lipid monolayer sealing the endosome, the resulting non-bilayer membrane rearrangement may expose the RNA — in complex with ionizable lipids — to the cytosol, without endosomal content release, but allowing endosomal escape of the RNA (5). In the neutral cytosolic environment, the ionizable lipids progressively deprotonate (6) and lose affinity for the RNA, which becomes increasingly hydrophilic and dissociates from the complex (7), yielding predominantly free RNA alongside a residual, transiently lipid-associated population. The model proposes sequence and molecular architecture, based on the experimental observations establishing cargo transfer, retention of the GUV pH gradient, interfacial remodelling in LUVs and heterogeneous diffusion of transferred siRNA.

This complements supported lipid bilayer studies (Aliakbarinodehi et al. 2024), allowing the testing of LNP components with respect to lipid re-arrangements and cargo translocation.

Our results distinguish cargo transfer from the membrane rupture assessed in cell-based studies by galectin recruitment (Munson et al. 2021). This distinction is critical for RNA therapeutics: if productive escape obligatorily required membrane rupture, increasing delivery would intrinsically promote a potentially harmful event, constraining efforts to improve efficacy without increasing damage-associated inflammation and toxicity. Preservation of membrane barrier function demonstrated by persistent pH gradient and absence of detectable open ruptures in LUV tomograms support siRNA transfer without membrane rupture or spatially restricted defects. Consistent with this distinction, Steinegger et al. reported that an MC3-based LNP formulation achieved efficient gene silencing despite low Galectin-8 recruitment, comparable to that of poorly delivering polyplexes. Their simulations nevertheless showed extensive membrane remodelling, including lipid exchange, leading to the proposal that limited membrane defects could reconcile effective delivery with weak damage readouts (Steinegger et al. 2026). These findings suggest that galectin recruitment does not consistently reflect productive escape across formulations. Endosomal damage may become more pronounced at high LNP exposure, raising the possibility that some damage-associated escape events reflect conditions exceeding those encountered at therapeutically relevant doses. The extent and kinetics of endosomal disruption also depend on lipid chemistry and exposure conditions and influence cytotoxicity (Bates et al. 2025). A recent preprint using GUVs and different ionizable lipid formulations reported cargo release associated with extensive membrane disruption (Pramanik et al. 2026). This study however did not resolve membrane barrier function combining measurements with cryo-electron tomography and characterisation of the transferred RNA. Our findings support an alternative explanation to obligatory rupture, in which siRNA translocation accompanies local lipid reorganisation without sustained loss of membrane barrier function. This provides a mechanistic basis for identifying formulations that enhance siRNA transfer while limiting membrane damage, with the prospect of separating delivery efficiency from damage-associated toxicity.

Protein exposure further modulated membrane transfer. Pre-incubation with serum or ApoE reduced luminal siRNA accumulation, consistent with reports of corona-mediated inhibition of delivery (Voke et al. 2025; Aliakbarinodehi et al. 2022). ApoE binding can redistribute LNP lipids and promote cargo release, (Sebastiani et al. 2021), suggesting that particle restructuring could contribute alongside steric shielding. Changes in surface lipid availability or cargo retention could consequently impair membrane-transfer activity. These observations highlight the need to balance protein-mediated cellular targeting with the subsequent capacity for cargo transfer.

The principal strength of our platform is its ability to separate experimentally accessible steps within a process that is difficult to dissect in cells. It distinguishes membrane association, lipid redistribution, cargo transfer and diffusion while independently controlling pH, lipid composition and protein exposure. The system does not reproduce the full molecular composition, leaflet asymmetry, geometry or dynamics of endosomes. Nevertheless, it makes individual steps experimentally accessible. The present findings concern an MC3-based formulation carrying siRNA but the approach can be extended to other lipids, cargos and membrane compositions (Pramanik et al. 2026). Coupling these measurements to cellular assays will help identify which properties can be predictive of functional delivery.

## MATERIAL AND METHODS

### Lipids and reagents

All phospholipids and cholesterol were purchased from Avanti Polar Lipids as powder or as stock solutions in chloroform unless otherwise stated. Phosphoethanolamine (cat. no. 852758), GM3 ganglioside (cat. no. 86005), phosphocholine (cat. no. 852467), cholesterol (cat. no. 700000), sphingomyelin (cat. no. 860061), DOPE (cat. no. 850725), DOPS (cat. no. 840035), PI3P (Echelon Biosciences, cat. no. P-3016), DOPC (cat. no. 850375), DiI and DiD (Invitrogen). SNARF-5F 5-(and-6)-carboxylic acid (Invitrogen, cat. no. S23922).

### LNP formulation

LNPs were formulated by microfluidic mixing using a NanoAssemblr Ignite (Cytiva Life Sciences) of a lipid phase in ethanol with an siRNA-containing aqueous phase (citrate buffer, 50 mM, pH 3), at an aqueous:organic flow rate ratio of 3:1 and a total flow rate of 10 mL/min. The lipid phase consisted of the ionizable lipid MC3, cholesterol, DSPC and PEG-DMPE at a molar ratio of 50:38.5:9.9:1.5 with 0.1 mol% DiD included for fluorescent labelling of the LNP lipid shell where indicated. The siRNA:ionizable lipid ratio was set to an N/P ratio of 5:1. Alexa488-labelled siRNA (siRNA-Alexa488) was used to enable fluorescence tracking of the RNA cargo. Following formulation, LNPs were diluted in 60 times their voume of PBS and then concentrated by centrifugal filtration (Amicon) to the desired final concentration.

### LNP characterization

LNP hydrodynamic diameter and polydispersity index (PDI) were determined by dynamic light scattering (DLS) on a Malvern Zetasizer. The formulation characteristics are reported in Table S1. siRNA-488 concentration in LNPs was quantified using a plate-reader-based fluorescence assay exploiting the Alexa488 label on the siRNA: LNP samples were measured after lysis with 0.1% v/v Triton X-100, the latter disrupting the LNP and releasing all encapsulated siRNA to give the total fluorescence signal.

### GUV preparation

Giant unilamellar vesicles (GUVs) were prepared by PVA-assisted gel swelling. Coverslips (18 × 18 mm) were coated with 200 µL of 5% (w/w) polyvinyl alcohol (PVA, MW 146,000-186,000; prepared by slow dissolution in water at 90 °C under stirring) and dried on a hot plate for 40 min at 50 °C. An early endosome-like lipid mixture was used for all GUV experiments (Cezanne et al. 2020): phosphoethanolamine (12.9 mol%), GM3 ganglioside (9 mol%), phosphocholine (3.6 mol%), cholesterol (32.3 mol%), sphingomyelin (12.6 mol%), DOPE (6.8 mol%), DOPS (6.1 mol%), PI3P (1 mol%) and DOPC (15.65 mol%), together with the fluorescent membrane marker DiD or DiI (Invitrogen) at 0.05 mol%. The lipid mixture (5 mM total lipid concentration) was spread onto the dried PVA film (10 µL) and dried under high vacuum for at least 30 min to remove residual solvent. The lipid film was hydrated with 400 µL of swelling solution (280 mM sucrose in 7 mM HEPES, pH 7.2) for 30 min at 48°C and then allowed to cool to room temperature. GUVs were gently detached from the PVA substrate by pipetting with a cut pipette tip and transferred to a fresh tube.

### LNP-GUV interaction assay

GUVs were mixed with siRNA-Alexa488-loaded LNPs (naked or preincubated with FBS or ApoE) diluted in imaging solution (280 mM glucose in 7 mM HEPES, pH 7.2), and a pH-switch buffer (140 mM sucrose + 140 mM glucose in 7 mM HEPES, freshly prepared and acidified for each experiment) was added to trigger LNP-membrane interaction. The volume of pH-switch buffer required to reach a final pH of 6.5 was determined for each batch in a test mixture, and the volume of imaging solution was adjusted to maintain constant final LNP and siRNA concentrations. Control samples received buffer at pH 7.2 to maintain neutral pH. Mixtures were incubated for 10 min at room temperature before imaging on a Zeiss LSM 780 confocal laser scanning microscope (inverted Axio Observer.Z1 stand), with 40x water-immersion objective, unless otherwise indicated (e.g. 60 min for extended-incubation experiments, Fig. 2).

### Luminal pH measurements

To monitor the pH inside and outside GUVs, the ratiometric pH indicator SNARF-5F 5-(and-6)-carboxylic acid (2.5 mM stock) was added at a 1:1000 dilution to the swelling solution used for GUV formation, so that the dye was encapsulated in the GUV lumen, and to the pH-switch buffer and imaging solution, so that it was also present, at a similar concentration, in the external medium. The swelling solution additionally contained 10 mM NaCl to provide a measurable ionic exchange across the membrane upon acidification in presence of gramicidin. GUVs and sample mixtures were prepared as described above. For each condition (pH 7.2 or 6.5, with or without LNPs and gramicidin), 10 large fields of view were acquired per sample, on a Zeiss LSM 780 confocal laser scanning microscope (inverted Axio Observer.Z1 stand), with a 40x water-immersion objective, at 10 min and 60 min after LNP or Gramicidin addition. SNARF-5F was excited at 488 nm, and emission was collected in two channels (560-600 nm and 620-660 nm), with the pinhole set to 1 Airy Unit; laser power, gain and pinhole were kept identical to the calibration acquisition. Individual GUVs and the surrounding bulk medium within each field were identified and measured separately during image analysis. For calibration, the ratio of the two SNARF channels was measured in sucrose buffers at 6 or more known pH values spanning the range about pH 6.0-8.0, containing SNARF at the same concentration as the samples, under identical imaging settings; the resulting ratio-versus-pH relationship was fitted with a linear regression to convert the ratio measured in the GUV lumen and in the external medium into pH. Calibration was repeated for each experimental session (Fig. S4). As a positive control for membrane permeabilization, gramicidin (final concentration 100 nM, from a freshly prepared ethanol stock) was added to GUVs at least 30 min before preparing the final mixture, in parallel with each condition.

### Serum and ApoE preincubation

LNPs were incubated with an equal volume of recombinant ApoE protein at 0, 5, 20 or 100 µg/mL (final ApoE concentration: half of the above), or with an equal volume of fetal bovine serum (FBS), for at least 1 h at 37 °C with shaking at 300 rpm, prior to addition to GUVs. For FBS-corona samples, unbound serum protein was removed by size-exclusion chromatography on an ÄKTA pure/start system (Cytiva) using a HiScreen™ Capto™ Core 400 column, to prevent the large molar excess of free serum protein from confounding membrane-interaction readouts through nonspecific protein-membrane adsorption. Naked LNPs (no FBS) were subjected to the same incubation and purification workflow, to control for any effect of the purification step itself. ApoE-corona samples were used without a purification step, as used at a defined, comparatively low total protein concentrations (2.5-50 µg/mL).

### LUVs for Cryo-electron microscopy preparation

Large unilamellar vesicles (LUVs) were prepared using the same lipid composition as the GUVs described above. A lipid film was generated at 5 mM total lipid concentration, dried under N₂ stream followed by vacuum for 1 h to remove residual solvent, and hydrated in 7 mM HEPES (pH 7.2) to a final concentration of 5 mM. The lipid suspension was incubated at 37 °C for 15 min and vortexed for 5 min, or until fully resuspended. The sample was then subjected to 10 freeze-thaw cycles and stored at −80 °C until use. LUV size distribution was checked by DLS as a quality control step prior to use (Table S1).

### Cryo-electron microscopy

Grids for electron microscopy (Multi A grids, Quantifoil) were glow-discharged and passivated with 0.1 mg/mL PLL(20 kDa)-g-PEG(2 kDa) (Alveole) in PBS for 30 min in a humidity chamber, then washed in Hepes buffer at pH 6.5 or 7.2. LNPs (in PBS), LUVs (in 7 mM HEPES, pH 7.2) and an acidic HEPES-based pH-switch buffer, freshly prepared and volume-adjusted for each batch to reach a final pH of 6.5 (or HEPES pH 7.2 to maintain neutral pH), were mixed and incubated for 10 min at room temperature before vitrification. A 3 µL aliquot of the mixture, supplemented with 0.75 µL of gold beads (BSA Tracer 10nm, Aurion), was applied to the grid and vitrified by plunge-freezing using a Leica GP1 (Chamber humidity = 95%, Chamber temperature = 22 degrees C, blot time = 2s, Ethane as cryogen). Cryo-electron tomography data were acquired on a Thermo Scientific Titan Krios G4 transmission electron microscope equipped with a Thermo Scientific Selectris X energy filter and a Thermo Scientific Falcon 4i direct electron detector, operated at 300 keV in zero-loss mode with an energy filter slit width of 10 eV. Tilt series were collected at underfocus (2-6 µm) at 64,000× magnification (0.195 nm pixel size at the specimen level) using SerialEM (Mastronarde 2005), with a dose-symmetric tilt scheme from −60° to +60° in 3° increments starting at 0°, and a cumulative electron dose of ∼120 e⁻/Å². Image stacks were motion-corrected and tomograms reconstructed using AreTomo (Zheng et al. 2022) or wartem/warp (Tegunov 2026). Tomograms shown in the figures were reconstructed at a pixel size of 1.56 nm and are displayed unfiltered/undenoised; XY slices shown have a thickness of 7.8 nm.

### Fluorescence correlation spectroscopy (FCS)

FCS measurements were performed by positioning the confocal volume in the middle of the GUV lumen, 20 µm from the coverslip, 10 min after LNP addition and acidification, on a Zeiss LSM 780 NLO confocal laser scanning microscope (inverted Axio Observer.Z1 stand), with Zeiss C-Apochromat W Korr UV-VIS-IR 40x/1.2. The confocal volume was obtained by using different concentrations of free Alexa Fluor 488 NHS-Ester (Succinimidylester) (Thermo Fisher Scientific, #A2000) in HEPES buffer (1-1000 nM) as a reference (Supplementary Figure S5a and b).

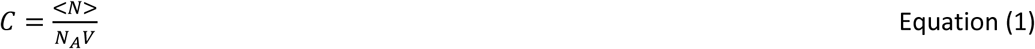

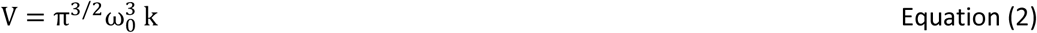

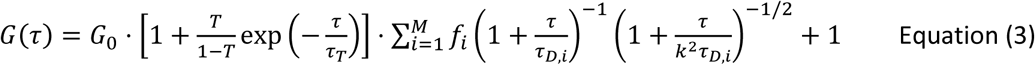

where C is concentration of Alexa488, <N> is average number of molecules in confocal volume, V. ω_0_ and k are the confocal width and structure parameter being the ratio of beam expansion in equatorial plane to axial plane. G(τ) is correlation function, where τ is the lag time for which correlation function is defined. T is the triplet fraction and *τ_T_* is corresponding triplet timescale. M is types of diffusing species, and *τ_D,i_* being the corresponding diffusion timescale and *f_i_* is fraction of i^th^ species. G_0_ is 1/<N>. Structure parameter is obtained from fitting single component with triplet to FCS curves. Fitting equation 1 and 2 to concentration vs average number of molecules <N> with already calculated structure parameter provides the confocal width. Using this confocal width, diffusion coefficient of free siRNA-Alexa488 (20nM, pH7.2, HEPES buffer) was determined (∼116 µm²/s at 23°C) (Hess and Webb 2002; Rüttinger et al. 2008). Further, in each experiment confocal volume was obtained using diffusion coefficient using equation (4):

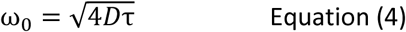

where τ is characteristic diffusion time of free siRNA-Alexa488 in HEPES buffer (pH 7, 20 nM) as a reference (1 AU = 32 μ*m*; laser power 7%; typical count rate ∼29-37 kHz, CPM ∼3.4-3.8). Measurements consisted of 10 repetitions of 10 seconds acquisitions per sample. Quickfit3.0 software was used to fit autocorrelation curves using multicomponent model and maximum entropy method (MEM) fit to obtain probability of diffusion timescales of species in a model independent method (Sengupta et al. 2003). MEM fits were analysed using custom script for predicting prominent timescales and calculating the diffusion coefficients. The detected timescales from single FCS were pooled together for further analysis. Outliers were removed by considering the characteristics timescale smaller than 40 µs (corresponding to photophysics of fluorphores) and applying generalized extreme studentized deviate (GESD) to remove further outliers. For experiments with FBS or ApoE pre-incubation, a multi-component fit was performed. For each recording, diffusion coefficients and molecule numbers were only retained for further analysis if the autocorrelation curve showed a clearly identifiable decay, the fit residuals showed no systematic pattern, the relative uncertainty (coefficient of variation) in the fitted molecule number (N) and diffusion coefficient (D) were both below 50%, and the fitted parameters were physically plausible. GUVs not meeting these criteria were excluded from the corresponding concentration and diffusion coefficient analyses, but were retained for count rate, which does not require autocorrelation fitting. This resulted in a smaller number of analysable GUVs for concentration and diffusion coefficient than for count rate, particularly for pre-incubated LNP samples, which typically yielded a weaker signal.

### Pre-formed siRNA-lipid complexes

siRNA-Alexa488 was diluted to 25 nM in 50 mM citrate buffer (CB), pH 3. MC3 and cholesterol working stocks were prepared by diluting the stock solutions in ethanol yielding working solutions of 1.246 mM (MC3) and 2.33 mM (cholesterol). Pre-formed siRNA-MC3 complexes (N/P 5) were generated by mixing MC3 working solution to the siRNA solution. The N/P ratio was calculated assuming 42 negative charges per siRNA duplex (21 bp) and full protonation of MC3 at pH 3, giving a target MC3 concentration of 5.25 µM. siRNA-MC3-cholesterol complexes were generated by mixing the MC3 and the cholesterol working solutions with the siRNA solution; the cholesterol concentration (4.04 µM) was set to match the cholesterol:MC3 molar ratio of the LNP formulation (38.5:50 mol%, ratio 0.77). Free siRNA controls were prepared in parallel in HEPES (7 mM, pH 7.2), in GUV imaging buffer (pH 7.2), and in CB (pH 3, with same volume of ethanol added as a vehicle control matching the ethanol content of the lipid-containing samples). All lipid-containing samples were vortexed for 5-10 s and incubated for 10 min before measurement. To assess the pH-dependence and reversibility of RNA-lipid association, aliquots of the pH 3 samples (free siRNA, siRNA-MC3 and siRNA-MC3-cholesterol) were sequentially titrated to pH 6.5 and then to pH 7.2 using NaOH. Samples were incubated for 10 min after reaching the target pH before FCS measurement. For all FCS measurements, laser power, pinhole and gain were kept identical across all samples within a session (See the above Fluorescence correlation spectroscopy (FCS) section).

### Statistics

Statistical tests, sample sizes (n, number of GUVs or measurements) and the number of independent experiments (N) are reported in the corresponding figure legend for each experiment. Unless otherwise stated, data are shown as mean ± s.e.m. of per-experiment medians across independent experiments, with statistical comparisons performed on these per-experiment values. Statistical analyses were performed in GraphPad Prism (version 11.1.0, GraphPad Software).

