## Supporting Information for "Reconstitution of lipid nanoparticle-mediated siRNA transfer across endosomal membrane mimics"

**Table S1. Hydrodynamic size and polydispersity of LNP and LUV preparations used in this study.** z-average diameter and polydispersity index (PDI) were measured by dynamic light scattering (Zetasizer, see Methods) for each formulation used across the figures indicated. Data are mean ± s.d. from N ≥ 3 independent batches.

| Formulation | Cargo | Label | Used in | Z-average (nm) (±SD) | PDI (±SD) |
|---|---|---|---|---|---|
| LNPs (MC3:DSPC:Chol:DMPE-PEG - 50:38.5:9.9:1.5) | Scrambled siRNA-Alexa488 | siRNA-Alexa488 | Fig. 4, 5 (FCS) | 92 (±20) | 0.13 (±0.07) |
| LNPs (MC3:DSPC:Chol:DMPE-PEG - 50:38.5:9.9:1.5) | Scrambled siRNA-Alexa488 | siRNA-Alexa488 + DiD (dual-labelled) | Fig. 1 and S2 | 105 (±10) | 0.14 (±0.10) |
| LNPs (MC3:DSPC:Chol:DMPE-PEG - 50:38.5:9.9:1.5) | anti-GFP siRNA | - | Fig. 2 and 3 | 114 (±16) | 0.12 (±0.06) |
| LUVs (early endosome-like) | - | - | Fig. 3 | 215 (±30) | 0.19 (±0.03) |

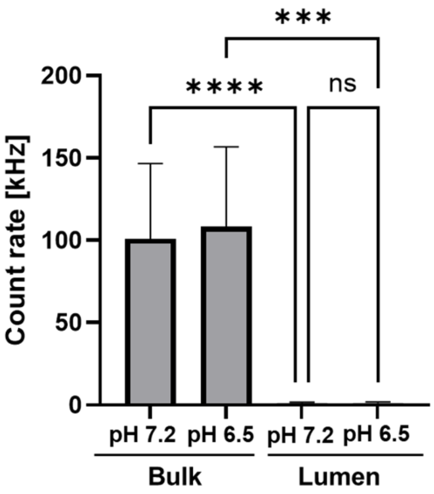


**Figure S1. Free siRNA does not enter the GUV lumen, independent of pH.** Corresponding to the confocal images in Fig. 1d, photon count rate was measured from a confocal detection volume (the same point-detection setup used for FCS measurements, see Fig. 3 and M&M) 10 min after GUV + free siRNA incubation and pH adjustment, in the bulk solution surrounding the GUVs (n = 3–10 measurements per condition) and inside the GUV lumen (n = 3–10 GUVs per condition), from N = 4 independent experiments. Data are shown in linear scale (kHz); statistical comparisons were performed by repeated-measures one-way ANOVA assuming a lognormal distribution, with Tukey's multiple comparisons test. ****P < 0.0001; ***P < 0.001; ns, not significant. Only comparisons relevant to assessing luminal translocation are shown. Bulk count rate confirms that free siRNA is present and detectable in solution at both pH values, while the luminal count rate remains at background level, confirming that free siRNA does not spontaneously cross or associate with the GUV membrane, regardless of external pH.

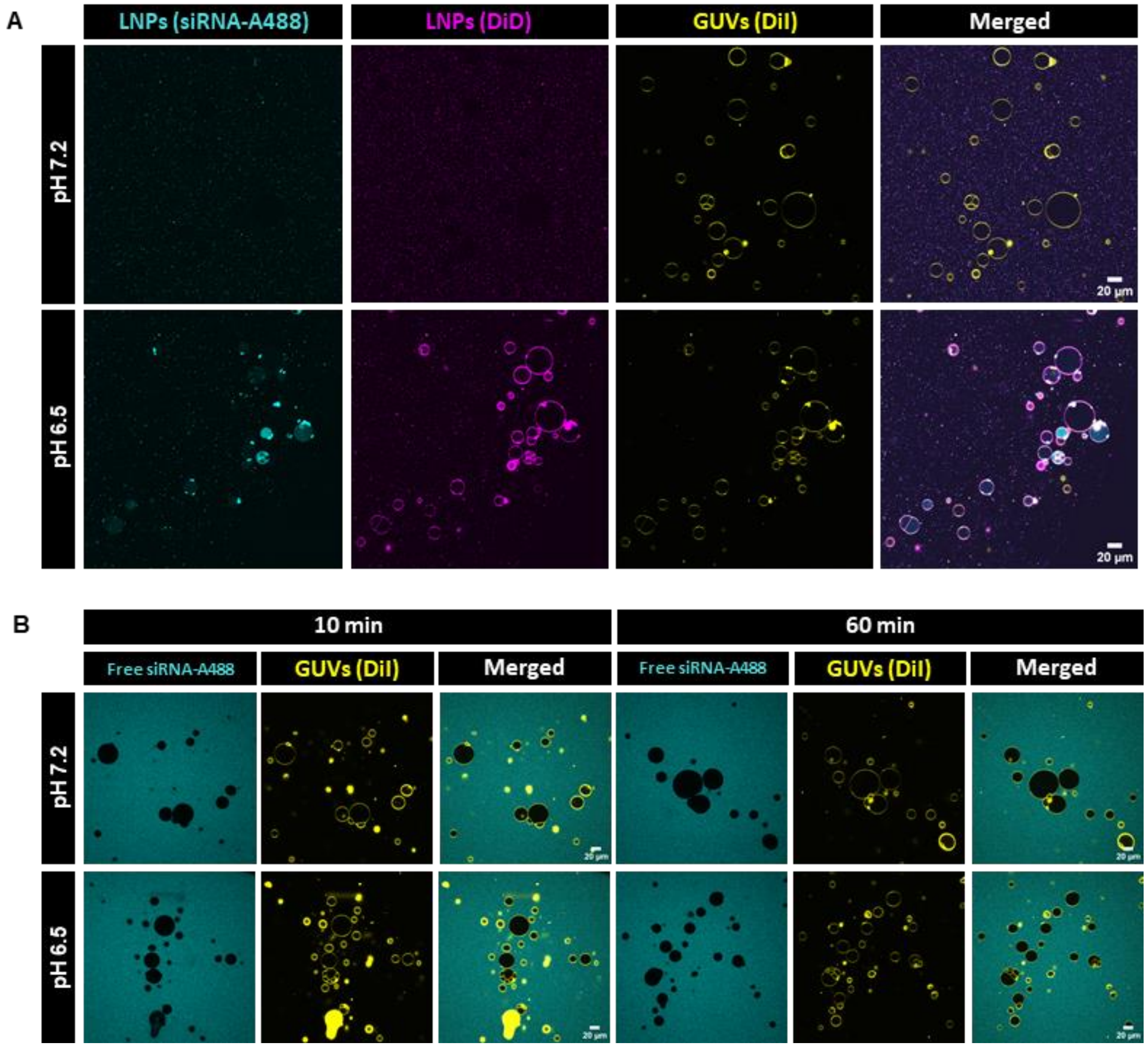


**Figure S2. Large-field-of-view images of LNP–membrane interaction and free siRNA controls. a,** Representative large-field-of-view confocal images of GUVs incubated for 10 min with dual-labelled LNPs (siRNA-Alexa488, cyan; DiD, magenta) at an external pH of 7.2 (top) or 6.5 (bottom), with the GUV membrane labelled with DiI (yellow), shown to illustrate the full imaged field. **b,** Representative large-field-of-view confocal images of GUVs incubated with free siRNA-Alexa488 (no LNPs) at pH 7.2 (top) or 6.5 (bottom), for 10 min or 60 min, shown for the same purpose. Scale bars, 20 µm.

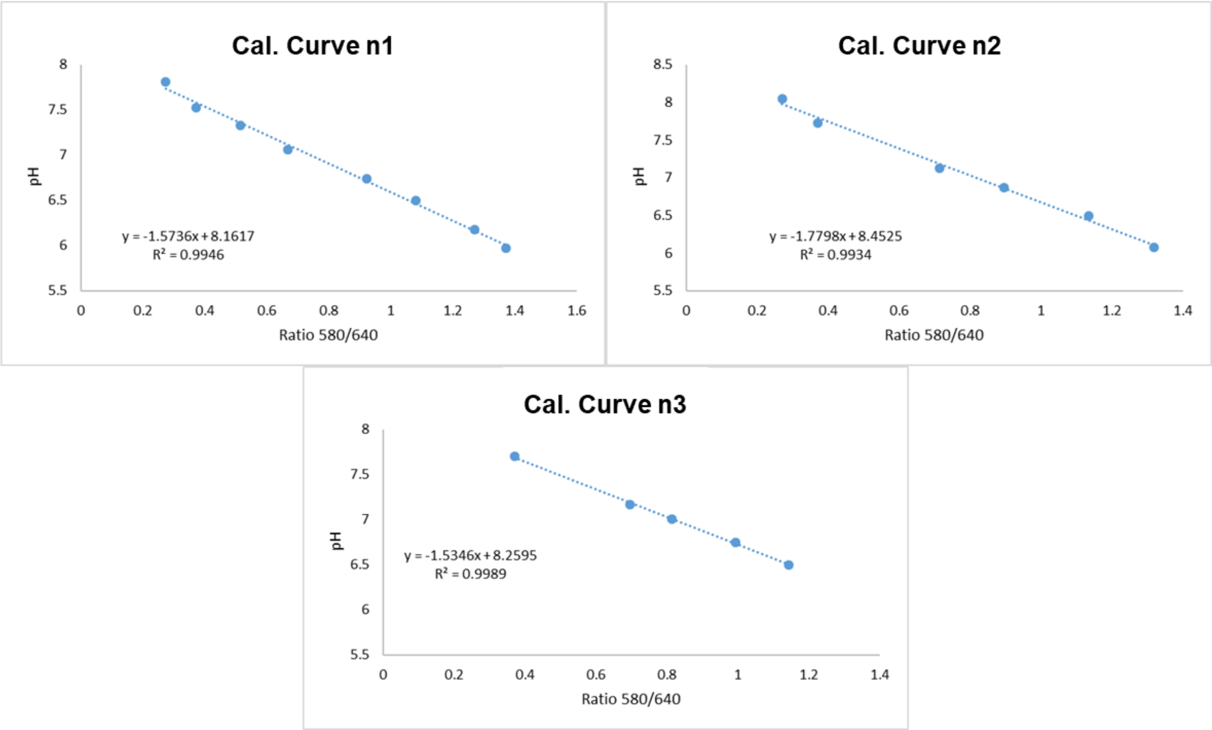


**Figure S3. SNARF-5F ratiometric calibration curve.** Ratio of the two SNARF-5F emission channels (560–600 / 620–660 nm) measured in sucrose buffers at known pH values, for two independent calibration sessions (Day 1: $y = -1.57x + 8.16$, $R^2 = 0.995$; Day 2: $y = -1.78x + 8.45$, $R^2 = 0.993$; Day 3: $y = -1.53x + 8.25$, $R^2 = 0.998$), each fitted by linear regression. Calibration was repeated for each experimental session (see Methods) to account for day-to-day variation in instrument sensitivity, as illustrated by the modest difference between the two curves shown.

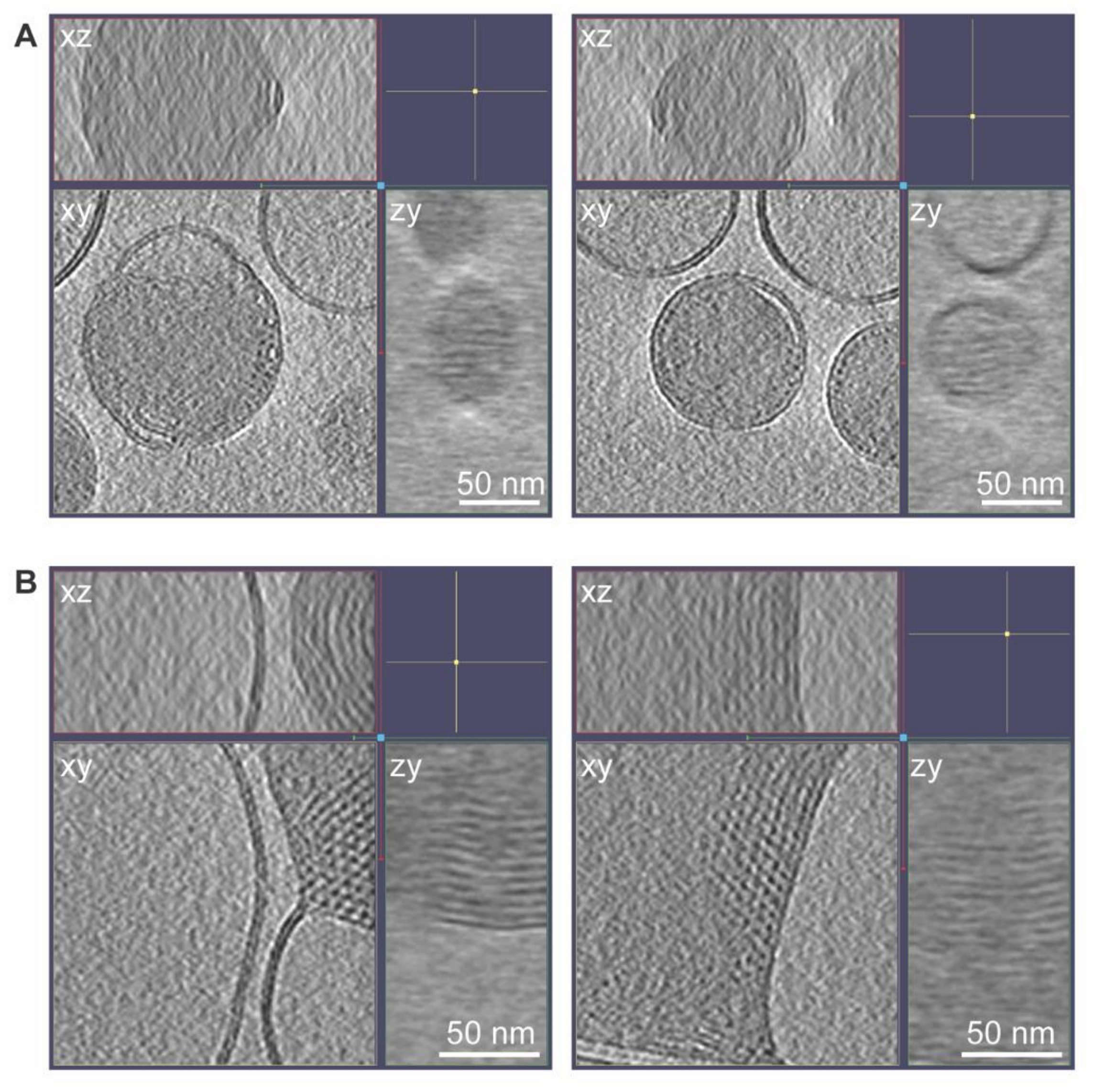


**Figure S4. Orthogonal views of ordered lipid arrangements in intact LNPs at neutral pH and in LNPs melted into LUV membranes at low pH.** XYZ views of two intact LNPs at neutral pH (**a**) and of two LNP assemblies melted into LUV membranes at low pH (**b**). Each window shows an XY plane (lower left), a YZ plane (lower right) and an XZ plane (upper left). The crosshair in the XY plane marks the point where the planes intersect. Electron-dense puncta in the XY plane are cross-sections of lines, as seen in the XZ and YZ planes. Scale bars, 50 nm.

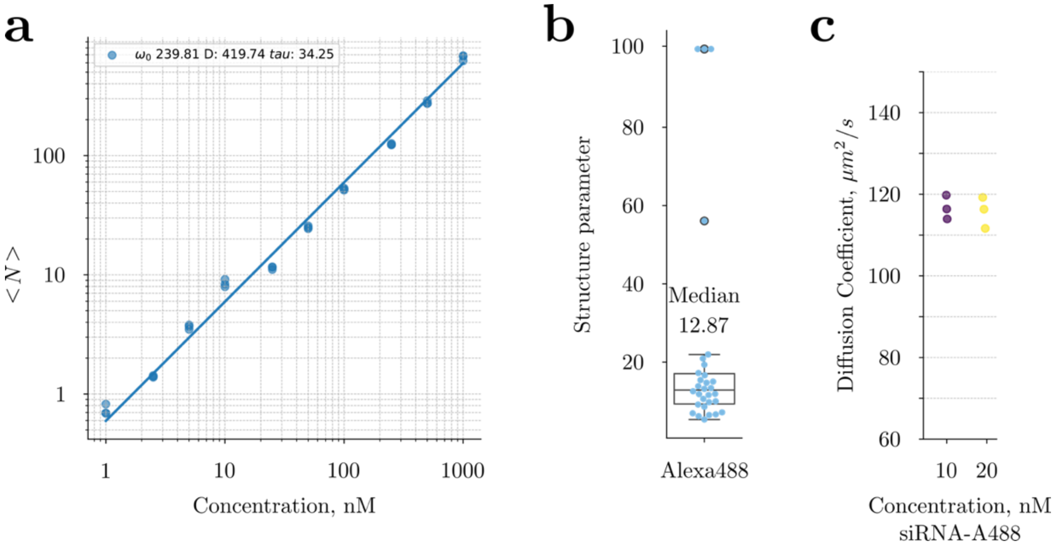


**Figure S5. FCS calibration and validation of concentration measurements. a**, Concentration calibration curve: number of molecules in the confocal volume (⟨N⟩) plotted against known Alexa488 concentration (log-log scale), fitted to obtain the beam waist ($\omega_0$ = 239.81 nm, close to the expected value of 244.34 nm) and diffusion coefficient (D = 419.74 $\mu m^2/s$, close to the expected 435.739 $\mu m^2/s$) of Alexa488. **b**, Distribution of the fitted structure parameter (axial-to-radial ratio of the confocal volume) across individual measurements (median = 12.87), used to constrain subsequent fits. **c**, Diffusion coefficient ($\mu m^2/s$) of free siRNA-Alexa488 obtained from the calibration in **a** and **b**, for two independent known concentrations (10 and 20 nM, purple and yellow respectively).
